# Longitudinal dentate nuclei iron concentration and atrophy in Friedreich ataxia: IMAGE-FRDA

**DOI:** 10.1101/464537

**Authors:** Phillip G. D. Ward, Ian H Harding, Thomas G. Close, Louise A Corben, Martin B Delatycki, Elsdon Storey, Nellie Georgiou-Karistianis, Gary F Egan

**Affiliations:** Monash Biomedical Imaging, Monash University, Melbourne, Australia.; School of Psychological Sciences and Monash Institute of Cognitive and Clinical Neurosciences, Monash University, Melbourne, Australia.; Australian Research Council Centre of Excellence for Integrative Brain Function, Melbourne, Australia.; Bruce Lefroy Centre for Genetic Health Research, Murdoch Childrens Research Institute, Melbourne, Australia; Department of Paediatrics, University of Melbourne, Parkville, Australia.; Victorian Clinical Genetics Service, Parkville, Australia.; Department of Medicine, Monash University, Melbourne, Australia

## Abstract

**Background:** Friedreich ataxia is a recessively inherited, progressive neurological disease characterised by impaired mitochondrial iron metabolism. The dentate nuclei of the cerebellum are characteristic sites of neurodegeneration in the disease, but little is known of the longitudinal progression of pathology in these structures.

**Methods:** Using *in vivo* magnetic resonance imaging, including quantitative susceptibility mapping, we investigated changes in iron concentration and volume in the dentate nuclei in individuals with Friedreich ataxia (n=20) and healthy controls (n=18) over a two-year period.

**Results:** The longitudinal rate of iron concentration was significantly elevated bilaterally in participants with Friedreich ataxia relative to healthy controls. Atrophy rates did not differ significantly between groups. Change in iron concentration and atrophy both correlated with baseline disease severity or duration, indicating sensitivity of these measures to disease stage. Moreover, atrophy was maximal in individuals early in the disease course, while the rate of iron concentration increased with disease progression.

**Conclusions:** Progressive dentate nuclei pathology is evident *in vivo* in Friedreich ataxia, and the rates of change of iron concentration and atrophy in these structures are sensitive to the disease stage. The findings are consistent with an increased rate of iron concentration and atrophy early in the disease, followed by iron accumulation and stable volume in later stages. This pattern suggests that iron dysregulation persists after loss of the vulnerable neurons in the dentate. The significant changes observed over a two-year period highlights the utility of quantitative susceptibility mapping as a longitudinal biomarker and staging tool.

## Introduction

Friedreich ataxia (FRDA) is the most prevalent inherited ataxia^1^. The neurological component of FRDA is characterised by progressive loss of motor coordination. In 96% of affected individuals, it is due to a homozygous GAA trinucleotide repeat expansion in intron 1 of the *FXN* gene, which leads to transcriptional repression of frataxin^2^. Frataxin is an essential mitochondrial protein, with loss resulting in impaired cellular iron homeostasis, disrupted energy metabolism and oxidative stress, which leads to cell death in affected tissues^3^. Iron was first implicated in the pathogenesis of FRDA when marked increase in mitochondrial iron was identified in a yeast frataxin knockout^4,5^. This led to the recognition of the importance of earlier work where iron deposits in myocardium from individuals with FRDA was identified^6^. Increased mitochondrial iron was then identified in fibroblasts from people with FRDA^7^.

In the central nervous system, pathology preferentially targets the dentate nuclei of the cerebellum, as well as the dorsal root ganglia and dorsal tracts of the spinal cord. Functional and structural changes are also evident in the cerebellar cortex, cerebello-cerebral white matter pathways, subcortical grey matter, and selectively in the cerebrum^8–11^.

Iron dysregulation is central to the mitochondrial metabolic deficits and subsequent neuropathology that underlies FRDA^12^. In the dentate nuclei, post-mortem histology has revealed significant atrophy of the large glutamatergic neurons, potentially secondary to gliosis, and redistribution of iron within the structure^12^. *In vivo* magnetic resonance imaging (MRI) studies have replicated findings of dentate atrophy at the macroscopic level, and studies using MRI relaxometry techniques have reported findings consistent with increased iron concentration^13,14^. More recently, Harding et al.^15^ corroborated these findings using quantitative susceptibility mapping (QSM), an emerging MRI technique, that provides a measure of heavy metal loading in the brain by measuring magnetic susceptibility^16^. QSM has excellent contrast at the borders of iron-rich structures that allows for accurate volume quantification. Harding et al. found that both dentate volume and QSM correlated with disease severity, and higher QSM was associated with lesser volume^15^.

Published cross-sectional observational studies suggest that iron loading and dentate nucleus integrity may be key markers of disease progression in FRDA, but longitudinal examinations have been lacking. One available study from Bonilha da Silva et al.^13^ reported reduction in T2 relaxometry – indicative of iron concentration increases – over a one year period. Further longitudinal characterisation of the natural history of dentate nucleus and broader central nervous system disruptions in FRDA is necessary to define the *in vivo* pathological course of the disease and address the current lack of validated biomarkers of disease progression for use in FRDA trials.

This study aimed firstly to assess longitudinal changes in iron concentration and volume in the dentate nuclei over a two-year period using QSM in individuals with FRDA relative to control participants. Secondly, we investigated whether the longitudinal effects were associated with changes in the behavioural and clinical measures, and whether there was evidence for staging effects whereby the trajectory of the changes depended on the stage of the disease. We hypothesised that the change in QSM, and of atrophy of the dentate nuclei, would be greater in individuals with FRDA than in controls, and that these changes would be associated with increased severity of the disease.

## Methods and Materials

This study was approved by the Monash Health Human Research Ethics Committee, and participants provided written informed consent prior to undertaking the study.

### Participants

Data was acquired as part of the IMAGE-FRDA study^15^. Data for the longitudinal analysis was available for 20 FRDA and 18 control participants (Table 1). Of the 36 genetically-confirmed participants with FRDA and 37 age- and gender-matched controls recruited at baseline, 31 participants from each cohort returned for a two-year follow-up testing session (mean=2.01 years). A hardware storage failure resulted in intractable corruption of the QSM data from 10 FRDA and 11 control participants; one additional FRDA and two control participants were excluded due to significant motion artefacts or anatomical abnormalities.

**Table 1.** Participant demographics and clinical scores (mean ± standard-deviation).

|  | FRDA<br>(n=20) | Controls<br>(n=18) | Between group difference<br>(p-value) |
| --- | --- | --- | --- |
| Gender (F:M) | 8:12 | 9:9 | 0.55 |
| Age at baseline (years) | 34.3 ± 12.5 | 39.7 ± 13.3 | 0.21 |
| Age at follow-up (years) | 36.3 ± 12.5 | 41.7 ± 13.3 | 0.20 |
| FARS at baseline | 76.5 ± 28.6 | | |
| FARS at follow-up | 85.0 ± 28.3 | | |
| Age Onset (years) | 19.2 ± 9.1 | | |
| GAA1 | 558 ± 258 | | |
| GAA2 | 866 ± 242 | | |

No significant differences in gender or age were observed at baseline or follow-up (Table 1); however, a small difference was apparent in timing of the repeat scans (1.96 years between visits for individuals with FRDA and 2.07 years for controls, p=0.06). Age at time of measurement (baseline or follow-up), and time-between-scans were controlled for in all analyses. All participants were right-handed, excepting 1 individual in the FRDA cohort, as assessed using the Edinburgh Handedness Inventory. Disease severity was quantified at each time point using the Friedreich Ataxia Rating Scale (FARS)^17^ by an experienced clinician (MBD). The FARS was scored out of 167, with a greater score indicating greater disease severity. In addition, GAA repeat length in the shorter *FXN* allele (GAA1) and disease duration (time from symptom onset to present) was measured using published methodology. The shorter allele, GAA1, is responsible for the greater portion of frataxin expression^18^, and hence has a greater effect on disease severity.

### MRI Protocol

Whole-brain MRI scans were performed using a 3-Telsa Skyra (Siemens, Erlangen, Germany) with a 32-channel head-and-neck coil. The same scanning equipment and protocol was applied at baseline and follow-up, which included a T1-weighted magnetization-prepared rapid gradient-echo (MPRAGE) and a T2*-weighted dual-echo gradient-recalled echo (GRE). The parameters for the MPRAGE acquisition were: acquisition time=3.5minutes, repetition time=1900ms, echo time=2.19ms, flip-angle=9°, field-of-view=256×256mm^2^, voxels=1mm isotropic, 176 sagittal slices. The parameters for the GRE acquisition were: acquisition time=11.5mins, repetition time=30ms, echo times=7.38ms and 22.14ms, flip-angle=15°, field-of-view=230×230mm^2^, voxels=0.9mm isotropic, 160 axial slices.

### QSM Processing

Raw k-space data for the GRE acquisition was stored and reconstructed as per-coil, per-echo, real and imaginary volumes. A T2*-weighted magnitude image was constructed as the sum-of-squares of the GRE coil images. Bias was corrected using N4 from Advanced Normalisation Tools^19^ (ANTs). A brain mask was then calculated using FSL-BET^20^, and eroded using a spherical kernel with a 2-voxel radius.

Coil combination was performed as the complex sum of the Hermitian inner product of the two echoes, for each coil^21^. Phase and QSM processing was performed using STI-Suite v2.2 (https://people.eecs.berkeley.edu/∼chunlei.liu/software.html) with default parameters (unless specified). The phase of the combined image was unwrapped with Laplacian unwrapping^22,23^ using voxel padding of 12×12×12 and the background field was removed using V-SHARP^24,25^. QSM images were reconstructed using the iLSQR algorithm^26^ and referenced to the whole-brain mean. All image processing was performed within the Arcana neuroimaging framework^27^.

### Manual Dentate Nucleus Segmentation

The FRDA disease-related pathology in the cerebellum (evident as atrophy on imaging) precluded the use of automated registration and segmentation analysis tools. The dentate nuclei were therefore manually traced on the QSM images for both cohorts in two stages. Firstly, the left and right dentate nuclei were manually traced on the QSM images obtained at baseline by two observers blinded to disease stage. Secondly, the tracings were rigidly transformed to the follow-up images using FSL-FLIRT^28,29^ and two new observers, blinded both to disease stage and to whether the scans were acquired at baseline or follow-up, examined and edited the tracings. Magnetic susceptibility of the dentate nuclei was calculated as the mean QSM value within the manually traced mask following a 1-voxel morphological erosion. Dentate nuclei volume was calculated as the number of voxels within the traced mask multiplied by the voxel volume. Bilateral mean values were also calculated, as the mean of the left and right measurements in QSM and volume.

### Statistical Analysis

Group differences between individuals with FRDA and control participants were examined in the left and right dentate nucleus with measurements of QSM and volume. Longitudinal change (∆QSM and atrophy) was calculated as the difference between measurements at each visit, divided by the time in years between visits and thus presented as change-per-year throughout the paper. Healthy aging effects were estimated in the control participants and used to correct for age in both groups. Between-group differences in longitudinal changes were assessed using one-tailed two-sample t-tests. Correlations with disease severity at baseline (FARS score), change in disease severity (∆FARS score), time with motor symptoms (disease duration), and GAA1 were assessed using non-parametric Spearman’s rho in the participants with FRDA.

### Visualisation

*Post-hoc* visualisations of the dentate atrophy and QSM changes were undertaken in standard (MNI) space following statistical analysis of the traced dentate masks in participant space. To aid intuitive visualisation of disease stage effects (i.e., significant correlations with disease duration), the cohort was split into two even groups based on disease duration. Artefactual ∆QSM differences in the white-matter and near the brain surface were removed using a dilated dentate nucleus mask from the SUIT atlas^30^. Change in volume (atrophy) was calculated in native space before interpolation to MNI space. To do so, the baseline images were first rigid-body co-registered to the follow-up images in native space using FSL-FLIRT applied to the T2*-weighted magnitude images. For registration of the images from native space to MNI space, the whole-brain T2*-weighted magnitude image from each participant was rigid-body co-registered to their T1-weighted anatomical image at each time point using FSL-FLIRT. The T1-weighted anatomical images were then nonlinearly warped to MNI space using ANTs^31^. The rigid-body and non-linear warp were applied sequentially to interpolate the QSM images and tracings from each participants’ native space to MNI space.

## Results

### Between-Group Differences

Significantly greater longitudinal change in QSM (i.e., ∆QSM) was observed in participants with FRDA relative to the controls in both the left and the right dentate nucleus with medium to large effect sizes (Table 2). Individual participant level results are shown separately for the left and right dentate nucleus (Figure 1A, B). Inspection of the results in MNI space identified that the strongest effects were located towards the superior aspect of the left dentate nucleus (Figure 2B).

**Table 2.** Cross-sectional and longitudinal observations in QSM and volume of the dentate nuclei in FRDA and control participants (mean and standard-error), along with 95% confidence interval of mean difference, effect size (Cohen’s d) and p value of one-tailed, two-sample t-test. All data are corrected for age.

|  |  |  | FRDA | Controls | 95% CI | Cohen's d | p |
| --- | --- | --- | --- | --- | --- | --- | --- |
| QSM (ppb) | Baseline | L | 112.3 ± 4.7 | 83.7 ± 5.7 | [13.7, 43.5] | 1.3 | <10<sup>-3</sup> |
| | | R | 115.0 ± 6.4 | 89.3 ± 5.2 | [8.7, 42.6] | 1.0 | 0.002 |
| | Follow-up | L | 128.5 ± 4.8 | 90.2 ± 5.7 | [23.3, 53.3] | 1.7 | <10<sup>-5</sup> |
| | | R | 128.2 ± 6.8 | 89.4 ± 5.5 | [20.9, 56.8] | 1.4 | <10<sup>-4</sup> |
| | Longitudinal (Year<sup>-1</sup>) | L | 8.4 ± 2.1 | 3.3 ± 1.0 | [0.2, 9.9] | 0.7 | 0.02 |
| | | R | 6.7 ± 2.1 | 0.0 ± 1.0 | [1.8, 11.6] | 0.91 | 0.005 |
| Volume (mm<sup>3</sup>) | Baseline | L | 789 ± 38 | 1060 ± 50 | [-397, -146] | 1.4 | <10<sup>-4</sup> |
| | | R | 749 ± 33 | 1009 ± 57 | [-390, -130] | 1.3 | <10<sup>-3</sup> |
| | Follow-up | L | 691 ± 31 | 955 ± 49 | [-379, -148] | 1.5 | <10<sup>-4</sup> |
| | | R | 656 ± 30 | 934 ± 55 | [-403, -154] | 1.5 | <10<sup>-4</sup> |
| | Longitudinal (Year<sup>-1</sup>) | L | -46.1 ± 7.9 | -51.5 ± 8.1 | [-17.6, 28.4] | 0.15 | 0.32 |
| | | R | -51.7 ± 9.5 | -36.5 ± 9.0 | [-41.8, 11.4] | 0.38 | 0.13 |

**Figure 1.**
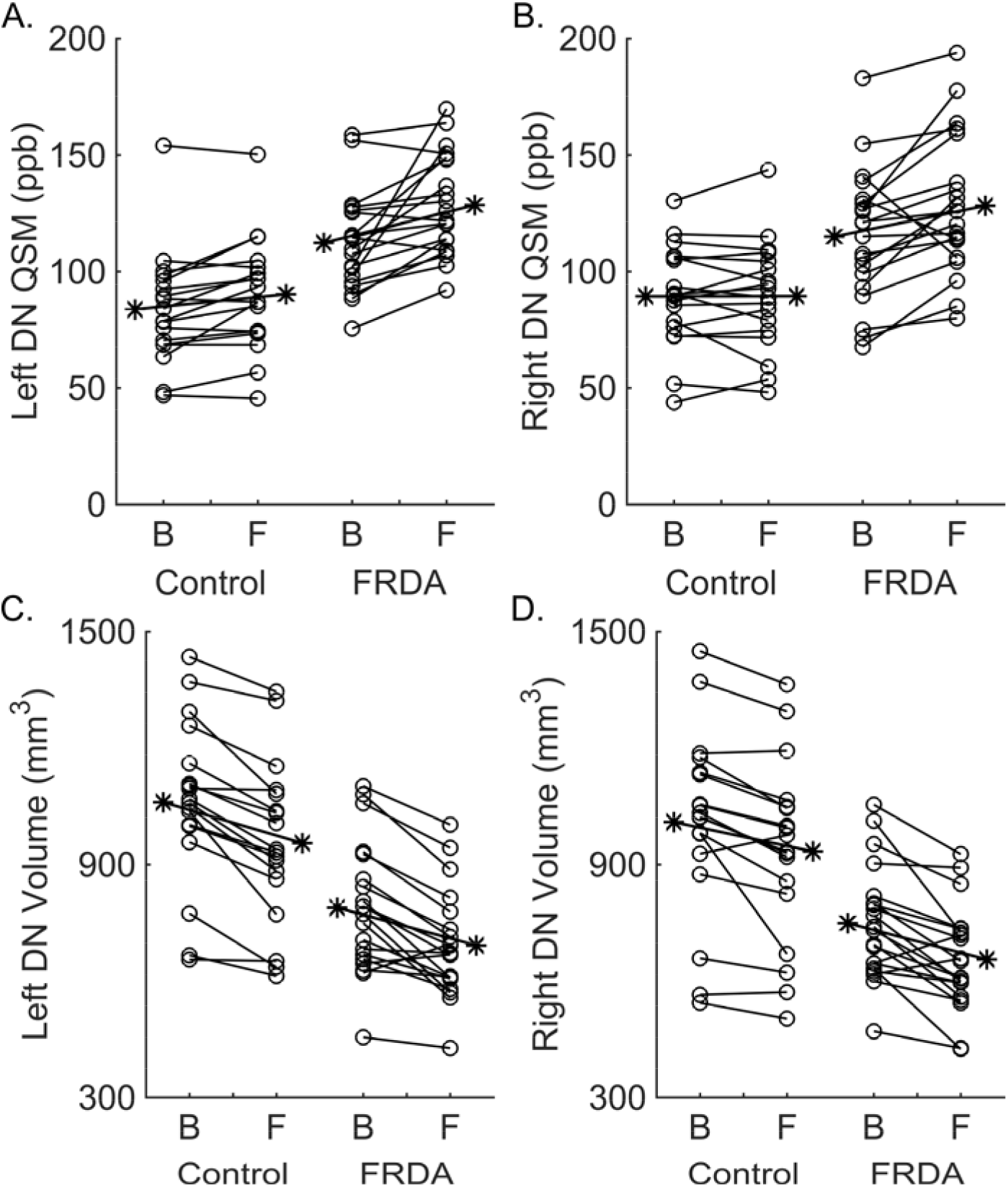
Mean QSM values (A, B) and volume (C, D) in the dentate nuclei (DN) for each participant in the FRDA and control cohorts at baseline (B) and follow-up (F). Values were controlled for age at each time-point independently. Group means are also overlaid as asterisks to the left and right of participant data for each group. ppb = parts per billion.

**Figure 2.**
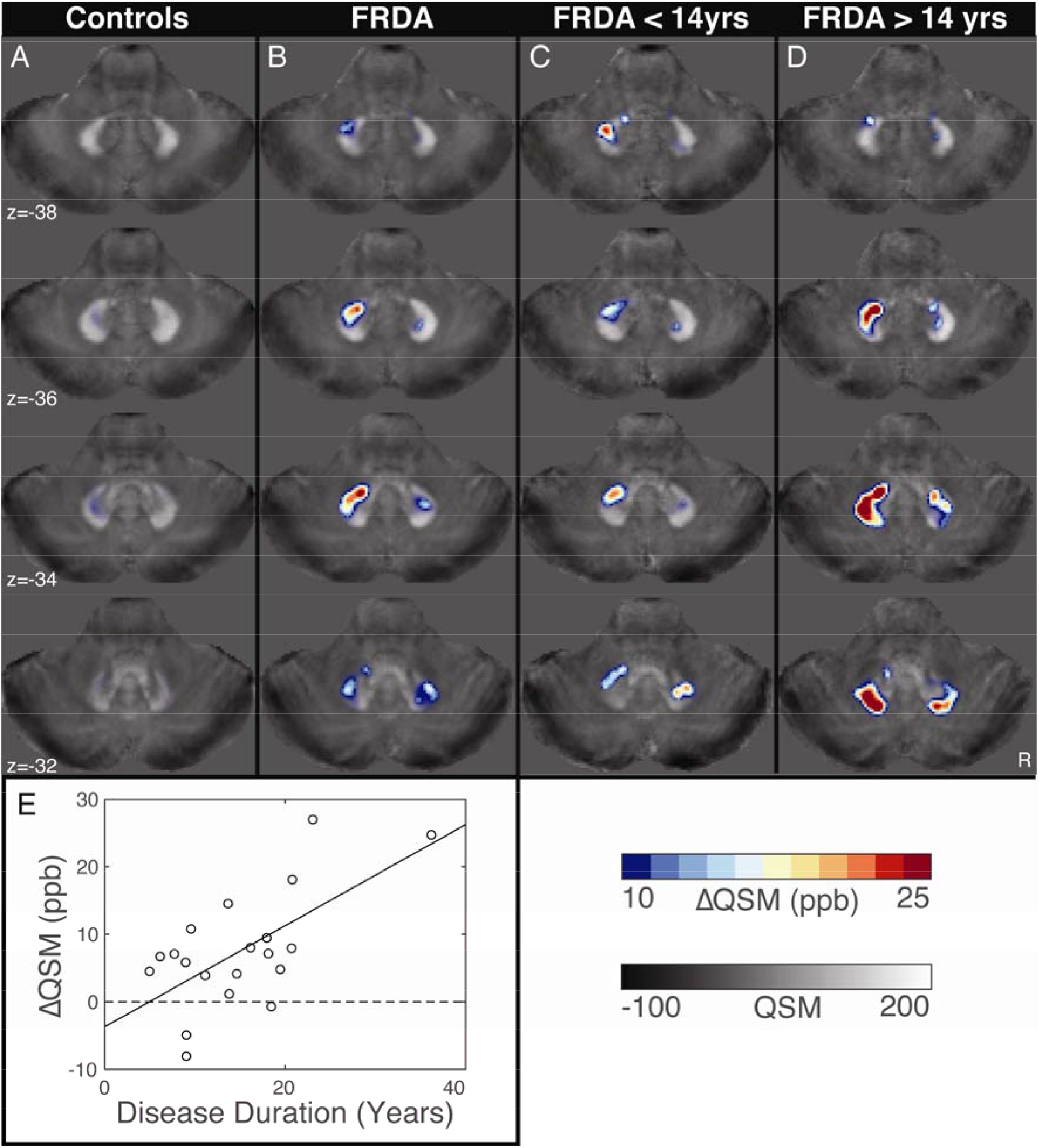
Longitudinal change in QSM (∆QSM) in participants from the control (A), and FRDA cohorts (B), and participants with less than, and greater than, 14 years of disease duration (C and D, respectively). Note that the median split of the sample (C, D) is undertaken only for visualisation, and all statistical inference is based on correlations across the full cohort. Changes are overlaid on group averages of QSM in MNI-space. Results are thresholded at 10 parts per billion (ppb) ∆QSM. (E) A linear correlation (solid line) between disease duration and ∆QSM, with a dotted-line to denote the average in the control group.

No significant between-group difference in the longitudinal rate of dentate nucleus atrophy was observed (Figure 3A, B; Table 2), although changes within both cohorts were pronounced (Figure 1C, D, Table 2).

**Figure 3.**
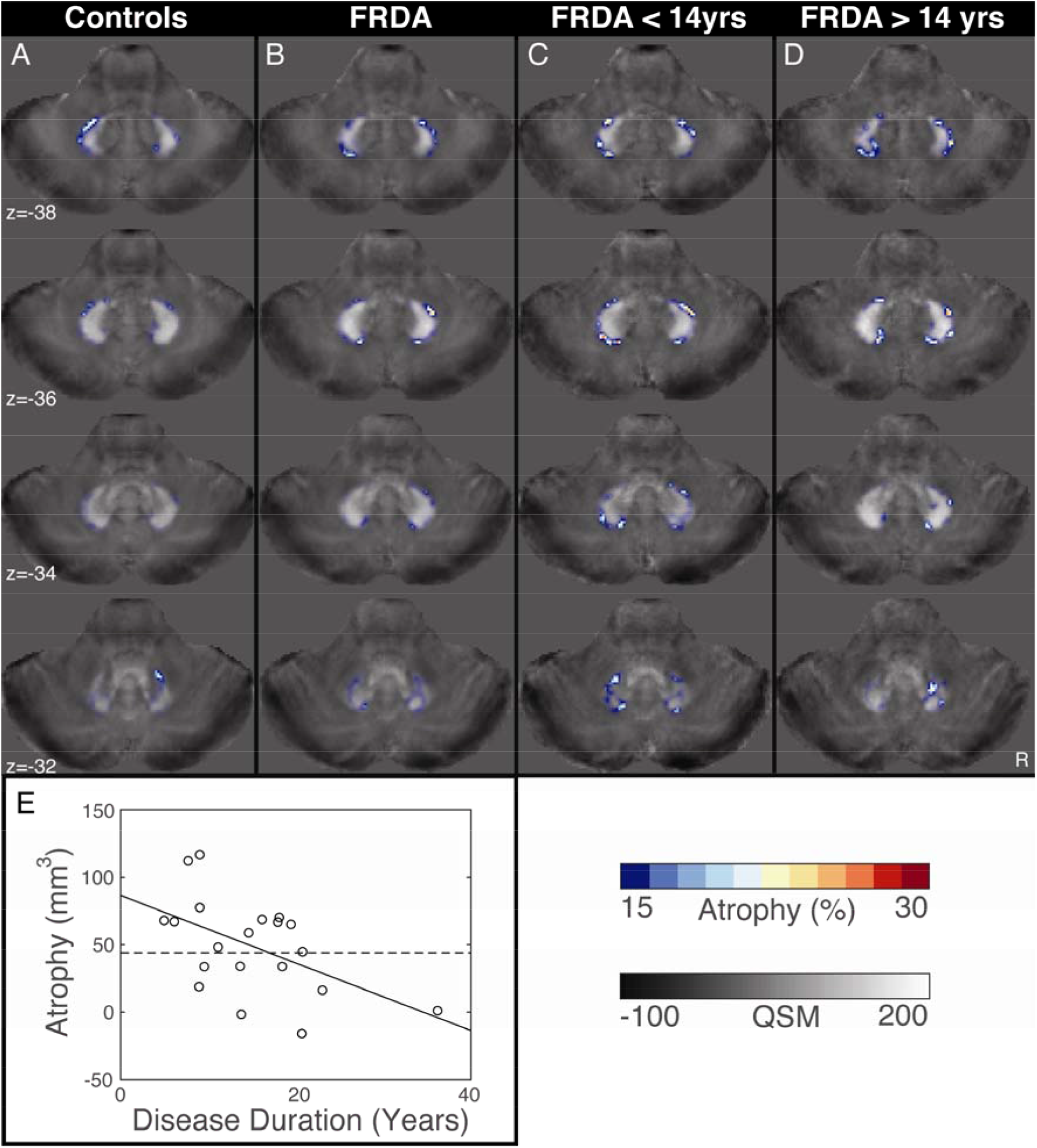
Longitudinal atrophy in participants from the control (A), and FRDA cohorts (B), and FRDA participants with less than, and greater than, 14 years of disease duration (C and D, respectively). Note that the median split is undertaken only for visualisation, and all statistical inference is based on correlations across the full cohort. Atrophy is shown in frequency, i.e., the percentage of participants who exhibited atrophy in a given voxel. Changes are overlaid on group averages of QSM in MNI-space. Results are thresholded at atrophy in 15% of participants. (E) A linear correlation (solid-line) between disease duration and atrophy, with a dotted-line to denote the average in the control group.

Cross-sectional differences in QSM and volume in the dentate nuclei were marked and showed consistency between the cerebellar hemispheres (Table 2). The confidence intervals of the group means were consistent over time for both QSM and volume.

### Clinical Correlations

Longitudinal ∆QSM values in the individuals with FRDA correlated bilaterally with disease duration (rho=0.49, p=0.03; Figure 2C, D, E), with lateralised trends (left: rho=0.40, p=0.08, right: rho=0.38, p=0.09). To account for differences in the severity of the pathophysiology across individuals, which may be confounding this apparent effect of disease progression, this relationship was confirmed while additionally controlling for GAA1 (bilateral: rho=0.60, p=0.008, left: rho=0.42 p=0.08, right: rho=0.56, p=0.02).

Despite no group differences in the rate of dentate nucleus atrophy, the rate of such atrophy in participants with FRDA was dependent on the disease duration. Disease duration correlated significantly with change in volume of the dentate nuclei (Figure 3C, D, E). The correlation was lateralised to the left cerebellar hemisphere (rho=0.66, p=0.002); right (rho=0.20, p=0.40), and persisted even after controlling for GAA1 and clinical severity (FARS score); rho=0.51, p=0.03.

The relationship between the disease stage and the rate of atrophy was also evidenced by a significant correlation between the atrophy rate and the FARS score at the time of the baseline scan (Spearman’s rho=0.56, p=0.01), indicating a greater rate of atrophy in individuals with less severe symptoms. The relationship was observed in both the left (rho=0.62, p=0.004) and the right dentate nucleus (rho=0.45, p=0.047).

GAA1 also trended negatively with atrophy in the right cerebellar hemisphere (rho=0.44, p=0.054). However, this relationship did not persist after controlling for disease severity (FARS) or duration (rho=0.20, p=0.44). No significant correlations were observed between changes in atrophy or QSM and disease severity (p=0.65 and p=0.88, respectively).

## Discussion

To our knowledge, this is the first study to investigate longitudinal changes in iron concentration and volume in the dentate nuclei using MR-based QSM in individuals with FRDA. Distinct trajectories of iron concentration and volume loss over a two-year period were observed. The rate of increase in iron concentration was found to accelerate with disease progression. Conversely, the rate of atrophy was highest early in disease expression and slowed to a plateau over time. This study indicates that key neuropathological changes underpinning the FRDA disease process occur at different stages and are observable *in vivo* longitudinally and cross-sectionally. Importantly, QSM-based biomarkers may be maximally sensitive to disease progression at different stages of the disease.

The dentate nuclei are known to be the most susceptible brain structures in the progression of FRDA pathology, with marked degeneration evident using both histological and neuroimaging approaches^15,32,33^. The present results indicate that early in the disease may be an especially vulnerable time, with atrophy in the dentate nuclei occurring maximally around the time of, or even prior to, onset of motor symptoms, and progressively decreasing over time. Interestingly, while this pattern was evident bilaterally with respect to disease severity (FARS score), a strong left-lateralised correlation between disease duration and rate of atrophy was also observed (which persisted even after controlling for disease severity). As all but one participant in this study was right-handed, this effect points to a particular vulnerability of the non-dominant motor hemisphere in early stages of motor symptom expression. It is interesting to note in this regard that one study has reported that function also degrades faster in the non-dominant limb in individuals with FRDA, with increasing preferential use of the dominant limb as the disease progresses^34^. Further detailed studies of bilateral motor function in individuals with FRDA would be required to corroborate this funding.

Iron dysregulation is considered central to the pathophysiology of FRDA^35^. Increased iron levels are associated with oxidative stress and neuroinflammation^36^, particularly in environments with high levels of non-transferrin-bound iron, as one would expect in an individual with a frataxin deficiency. Previous histological and cross-sectional neuroimaging studies have found evidence for a concentration or redistribution of paramagnetic substances (e.g., iron, copper, zinc) in the dentate nuclei of individuals with FRDA^12,33,37^. The present study indicates that changes in iron concentration in the dentate nuclei are progressive, corroborating a previous longitudinal study that used an alternative iron-sensitive neuroimaging approach (T2-relaxometry)^13^. This convergence of findings from studies using different MR based imaging techniques lends substantial support to the validity and utility of susceptibility-based imaging methods in non-invasively interrogating neuropathological changes in FRDA.

Importantly, changes in the QSM values in the dentate nuclei can be driven either by an absolute accumulation of iron or by a relative increase in iron concentration due to atrophy^38^. The noteworthy non-linear trajectories of the QSM and volume changes reported in this study suggest that both processes may be contributing to the changes observed in iron measurements at different disease stages. In individuals with more severe symptoms, an increased rate of change of the QSM values together with plateauing of the atrophy rates suggests greater iron accumulation in later stages of the disease. In contrast, more rapid progression of atrophy in the early stages of the disease was observed in parallel with slower change in the QSM values, possibly because the iron in the dentate nuclei was predominantly increasing in concentration rather than accumulating. These findings also indicate that iron dysregulation likely continues well after loss of the glutamatergic neurons in the dentate nuclei.

Notably, an elevation in the QSM values in the dentate nuclei could also occur through mechanisms independent of absolute iron increases, including redistribution of the concentrations of different iron species (free, ferritin-bound, heme-bound, sulphur-bound) or variation in concentrations of other bio-metals, such as copper^38^. This may reconcile apparent inconsistencies between the elevated QSM measurements reported in this and other *in vivo* studies, and histological studies that report redistribution of iron in the post mortem brain but do not observe increased iron quantity^12,33,37^. Thus, while our study provides evidence of progressive iron dysregulation in the dentate nuclei of individuals with FRDA, the precise mechanisms underlying changes in the *in vivo* QSM signal remains uncertain.

This study has several limitations. The FRDA cohort was restricted to adults who had a minimum disease duration of 5 years, and included five individuals (one-quarter of the sample) who were classified as “late-onset” (symptom onset >25 years old). Future work is necessary to extend these assessments of disease related effects and their variation in the dentate nuclei to minors as well as those in the early stage of disease.

Due to the relatively recent development of QSM techniques, there remains some contention around best practice for processing and intra-participant analyses^39,40^. In the current study, the whole brain mean susceptibility value was used as a reference value to avoid potential biases that could have arisen from unexpected disease-related susceptibility changes in the specific anatomical regions used in other studies^41–46^. However, the longitudinal changes in QSM within the control cohort in the present study were consistent with an increase in magnetic susceptibility as previously reported with age^47^. The cross-sectional values, in absolute terms, are also highly varied in the literature^41–46^, and do not conflict with those reported in this study. The differences in absolute quantification between published studies may be attributed to differences in QSM technique, tuneable parameters, phase processing methods, imaging protocol, scanner hardware and field strength. However, these factors do not impact the validity of between-group inference (relative quantification) within this or other studies.

## Conclusions

Quantitative magnetic susceptibility mapping is sensitive to the longitudinal progression of iron dysregulation in the dentate nuclei in individuals with FRDA. The QSM technique provides medium-to-large effect sizes of disease progression and is a promising biomarker for FRDA disease monitoring. Variable rates of pathological change in the dentate nuclei in FRDA were evident, with the early stage of the disease characterised by marked atrophy and moderate iron increases, followed by plateauing of atrophy and increasing iron levels. Further studies are required to investigate the complex interplay of atrophy and iron accumulation in the dentate nuclei, and to reveal the molecular mechanisms driving iron toxicity in FRDA.

## Acknowledgements

The authors thank the study participants for volunteering their time to the study. The authors also thank Monique Stagnitti, Richard McIntyre, and Parnesh Raniga for their assistance with recruitment, scanning, and data processing.

